# Paternal easiRNAs regulate parental genome dosage in Arabidopsis

**DOI:** 10.1101/203299

**Authors:** German Martinez, Philip Wolff, Zhenxing Wang, Jordi Moreno-Romero, Juan Santos-González, Lei Liu Conze, Christopher DeFraia, Keith Slotkin, Claudia Köhler

## Abstract

The regulation of parental genome dosage is of fundamental importance in animals and plants, exemplified by X chromosome inactivation and dosage compensation. The “triploid block” is a classical example of dosage regulation in plants that establishes a reproductive barrier between species differing in chromosome number ^1,2^. This barrier acts in the endosperm, an ephemeral tissue that nurtures the developing embryo and induces the abortion of hybrid seeds through a yet unknown mechanism. Interploidy hybridizations involving diploid (2×) maternal parents and tetraploid (4×) pollen donors cause failure in endosperm cellularization, leading to embryo arrest ^3^. Here we show that paternal epigenetically activated small interfering RNAs (easiRNAs) are responsible for the establishment of the triploid block-associated seed abortion in *Arabidopsis thaliana*. Paternal loss of the plant-specific RNA polymerase IV suppressed easiRNA formation and rescued triploid seeds by restoring small RNA-directed DNA methylation at transposable elements (TEs), correlating with reduced expression of paternally expressed imprinted genes (PEGs). We propose that excess of paternally derived easiRNAs in diploid pollen prevents establishment of DNA methylation, leading to triploid seed abortion. Our data further suggest that easiRNAs form a quantitative signal for chromosome number and their balanced dosage is required for post-fertilization genome stability and seed viability.

---

Previous work indicated that 24-nt small interfering RNAs (siRNAs) are sensitive to genome dosage and that their level is strongly reduced in response to interploidy hybridizations of 2× maternal parents and 4× pollen donors ^4^. To understand the genetic basis of this phenomenon, we tested the effect of several mutants impaired in the biogenesis of 21/22-nt and 24-nt siRNAs for their effect on establishing the triploid block. We generated double mutants with the *omission of second division* (*osdl*) mutant that forms unreduced (2n) male gametes ^5^. Thus, crossing *osd1* as pollen donor to wild-type (wt) plants results in triploid (3×) seed formation at almost 100% frequency ^5^, with the majority of 3× seeds being collapsed and non-viable ^6^ (Fig. 1a). Strikingly, using the double mutant *nrpd1a osd1* as pollen donor, viability of 3× seeds derived from those crosses was largely restored, with more than half of those seeds being non-collapsed and able to germinate (Fig. 1a,b). *NRPD1a* encodes for the largest subunit of RNA polymerase IV (Pol IV), a plant specific polymerase that transcribes heterochromatic regions of the Arabidopsis genome and leads to the production of 24 nt sRNAs that mediate DNA methylation ^7,8^. The effect of *NRPD1a* on triploid seed rescue was exclusively paternal, there was no significant rescue of triploid seed viability when *nrpd1a* plants were used as maternal parents in crosses with *osd1* and *nrpd1a osd1* (Fig. 1c). Triploid seed rescue by *nrpd1a* pollen was associated with restored endosperm cellularization (Fig. 1d), similar as observed for other mutants able to rescue triploid seed abortion ^6,9^. Together with *nrpd1a*, mutants in *DCL4*, *RDR6*, *AGO2*, and *AGO6* also significantly increased viable seed formation by a minimum of two-fold over background level, while no significant effect was observed for mutants in *DCL3*, *RDR2*, and *Pol V* (Fig. 1a). DCL2, DCL4, RDR6, AGO1, AGO2, and AGO6 are required for the production and function of 21/22nt easiRNAs targeting transcriptionally active TEs ^10–14^. In pollen, easiRNAs are produced in the vegetative cell and loaded into sperm cells ^14,15^; nevertheless, how they are produced and their downstream response remained to be identified. Given the strong effect of *nrpd1a* on triploid seed rescue, we addressed the question whether Pol IV and not Pol II could influence the production of 21/22-nt easiRNAs in pollen. We therefore sequenced sRNAs of pollen grains derived from 2× and 4× Coland *nrpd1a* mutant plants (1n and 2n Col and *nrpd1a* pollen, respectively, Fig. S1, Table S1). Pollen derived from 2× and 4× *nrpd1a* plants had not only strongly reduced levels of TE-derived 24-nt siRNAs, but as well strongly reduced levels of 21-nt and 22-nt siRNAs, pinpointing an undiscovered new role of Pol IV in the biogenesis of these sRNA classes in the male germline in plants (Fig. 2a, b). Analysis of the bias between sRNAs derived from the plus or minus strand of TEs indicated that indeed Pol IV is the major producer of dsRNA precursors for TE-derived sRNA biogenesis in pollen (Fig 2c). This effect is moreover exclusive to TEs and not to other epigenetically-regulated regions, like rRNA genes (Fig. 2d). However, in contrast to the dependency of Pol IV RNA biogenesis on RDR2 in somatic plant tissues ^16,17^, our genetic data do not support an exclusive role of RDR2 in this process in pollen, but rather implicate a role for RDR6 and possible redundancies with other RDRs, similar as suggested for diRNA synthesis ^18^. Consistently, production of 21/22-nt siRNA is reduced in *rdr6* and *rdr6 dcl4* double mutants (Fig. S2), supporting a role of DCL4 and RDR6 in the production of 21/22-nt siRNAs ^19–21^. Thus, our data revealed a new Pol IV-dependent pathway leading to the production of 21/22-nt and 24-nt siRNAs from TEs in pollen that potentially avoids the risk of forming potentially harmful full-length TE transcripts.

**Fig. 1.**
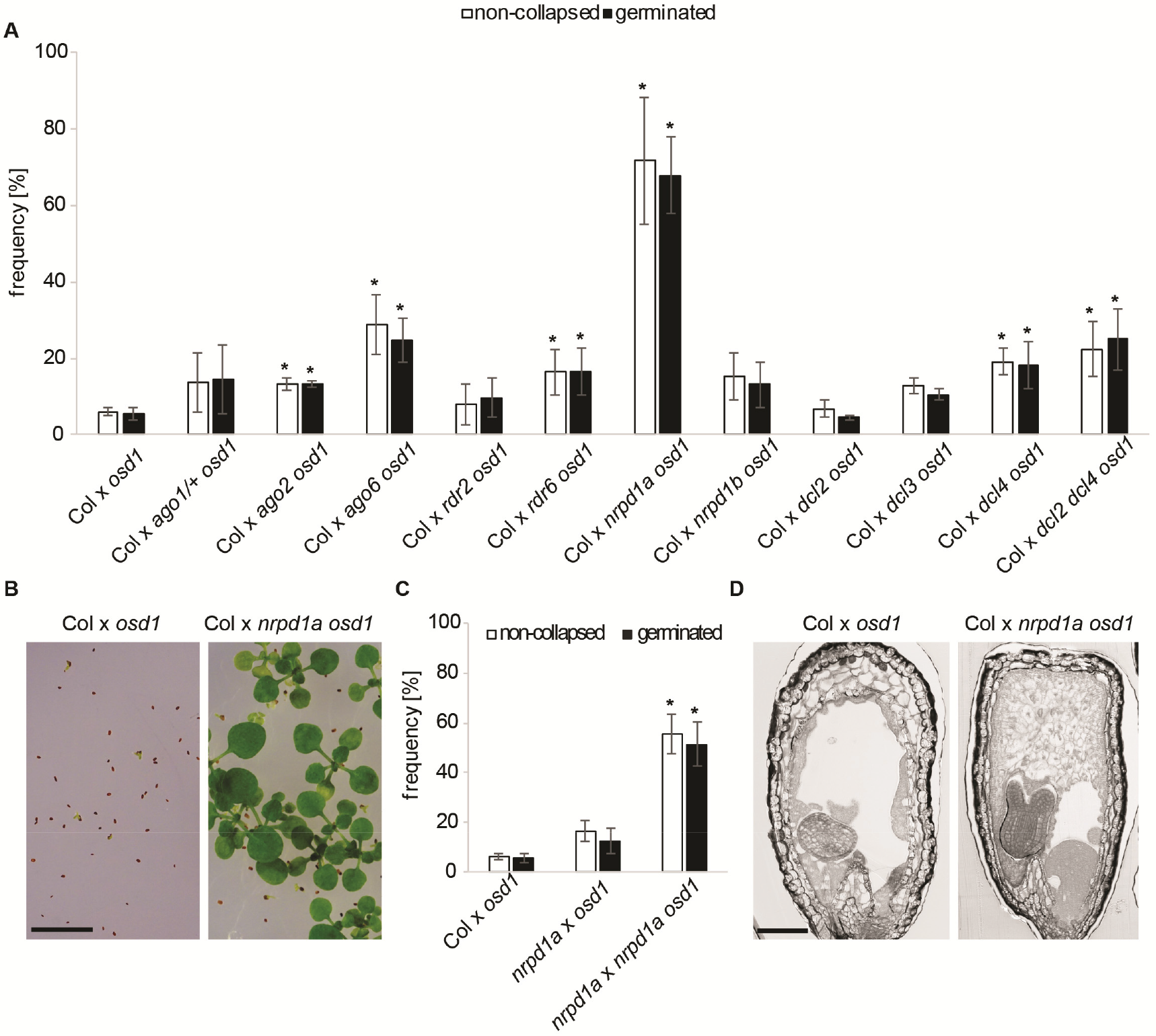
Pol IV mutant pollen is able to rescue seed abortion in triploid seeds. (**A**) Frequency of non-collapsed and germinated seeds derived from crosses of wild-type (Col) maternal parents with *osd1* and *osd1* double mutants of indicated genotypes. Asterisks mark significant differences (*P*<0.05) to the cross Col × *osd1* (Chi square test with Bonferroni correction). By convention, the female parent is always indicated first. (**B**) Pictures of non-germinating seeds (left panel) and seedlings (right panel) derived from the crosses Col × *osd1* and Col × *nrpd1a osd1*, respectively. Scale bar, 1 cm. (**C**) Analysis of the parent-of-origin effect of *nrpd1a* on triploid seed rescue. Asterisks mark significant differences (*P*<0.05) to the cross Col × *osd1* (Chi square test with Bonferroni correction). (**D**) Sections of 6 DAP seeds derived from Col × *osd1* and Col × *nrpd1a osd1* crosses. Scale bar, 0.1 mm.

We hypothesized that enlarged pollen grains of higher ploidy (Fig. S3) could potentially have an excess of 21/22-nt easiRNAs, building the triploid block. To test this, we purified RNA of 1n and 2n wt and *nrpd1a* mutant pollen samples normalized for an approximately equal number of pollen grains (Fig. 2e)^12^ and ribosomal RNA in 2n pollen RNA samples of wt and *nrpd1a* mutant plants compared to RNA samples of 1n pollen (Fig. 2f), strongly suggesting that there is a general increase of RNA in 2n pollen compared to 1n pollen.

**Fig. 2.**
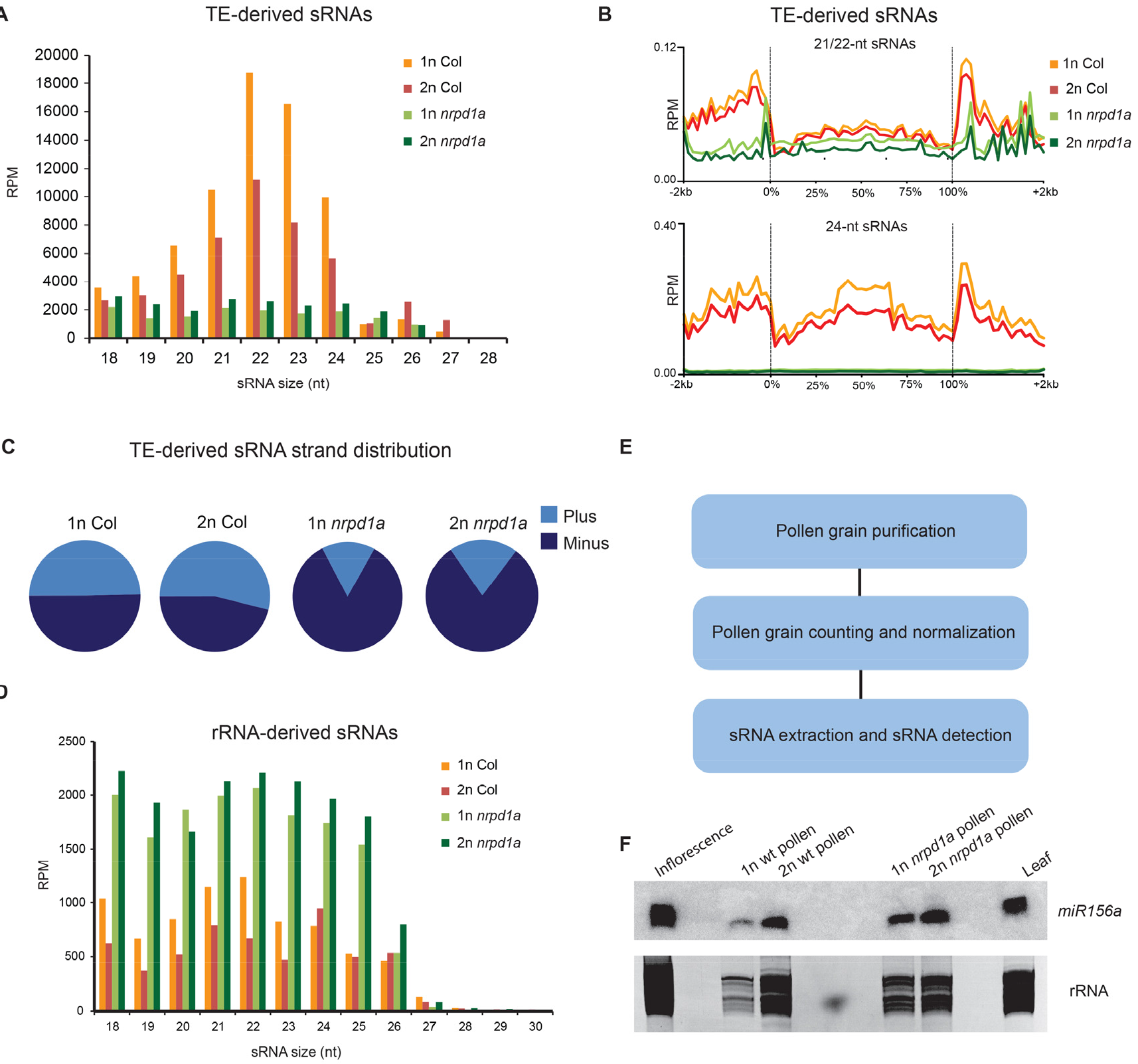
Pol IV is responsible for the biogenesis of pollen TE-derived sRNA in the size range of 19-24 nt. (**A**) sRNA profiles of TE-derived sRNAs of 1n pollen from Col wild-type (wt) and *nrpd1a* mutants and 2n pollen from tetraploid wt and *nrpd1a* mutants. (**B**) Metaplots of siRNA distribution over TE gene sequences in 1n and 2n pollen of Col wt and *nrpd1a* mutants. (**C**) sRNA strand origin percentage for TE-derived sRNAs in 1n and 2n pollen from Col wt and *nrpd1a* mutants. (**D**) sRNA profiles of ribosomal RNA (rRNA)-derived sRNAs of 1n and 2n pollen from Col wt and *nrpd1a* mutants. (**E**) Flowchart of steps for sRNA isolation used in the Northern blot shown in panel F. (**F**) Northern blot detection of *miR156a* in 1n and 2n pollen from Col wt and *nrpd1a* mutants. RNA samples were normalized to total amount of pollen grains. rRNA loading control from EtBr stained gel is shown in the lower panel.

Interploidy hybridizations using *osd1* pollen donors cause strong reduction of CHH methylation (H corresponds to A, T, or C) ^22^. Based on the previously proposed antagonistic relationship between post-transcriptional gene silencing (PTGS) mediated by 21/22-nt siRNAs and RNA-dependent DNA methylation (RdDM) ^12^, we raised the hypothesis that increased dosage of pollen-derived easiRNAs in 2n pollen negatively interferes with CHH methylation establishment in the endosperm. We tested this hypothesis by analyzing whether depletion of 21/22-nt easiRNAs from pollen restores CHH methylation in the endosperm. We analyzed DNA methylation in purified endosperm of 2× and 3× seeds derived from pollinations of L*er* maternal plants with wt Col or *nrpd1a* 1n or 2n pollen (referred to as 2× or 3× *nrpd1a* seeds, respectively, Fig. S4, Table S2). Consistent with previous work ^22^, CHH methylation at TEs was strongly reduced in the endosperm of 3× seeds and slightly increased in 3× *nrpd1a* seeds (Fig. 3a), indicating that loss of paternal PolIV function restores CHH methylation at defined TEs. About one quarter of TEs losing CHH methylation in 3× seeds restored CHH methylation in 3× *nrpd1a* seeds (Table S3, Fig. 3b, c). Strikingly, loci that experienced the strongest loss of CHH methylation in 3× seeds, had the highest gain of CHH methylation in 3× *nrpd1a* seeds (Fig. 3d), suggesting that easiRNAs negatively interfere with CHH methylation when present in increased dosage. Consistently, levels of 21/22-nt easiRNAs in pollen correlated with loss of CHH methylation in 3× seeds (Fig. 3e; Fig. S5a), suggesting that their increased dosage in 2n pollen causes the negative effect on CHH methylation establishment in 3× seeds. Loss of CHH methylation was PolIV dependent and correlated with loss of 21/22-nt easiRNAs in *nrpd1a* pollen (Fig. 3f, Fig. S5b) rather than with the presence of 24 nt siRNAs in 2n pollen (Fig 3h). Conversely, gain of CHH methylation in 3× *nrpd1a* seeds correlated with loss of 21/22-nt easiRNAs in pollen (Fig. 3g, Fig. S5c) and was not associated with reduced expression of *NRPD1a* in the endosperm (Fig. S6) Together, our data strongly suggest that pollen-derived 21/22-nt easiRNAs cause loss of CHH methylation in 3× seeds after fertilization.

**Fig. 3.**
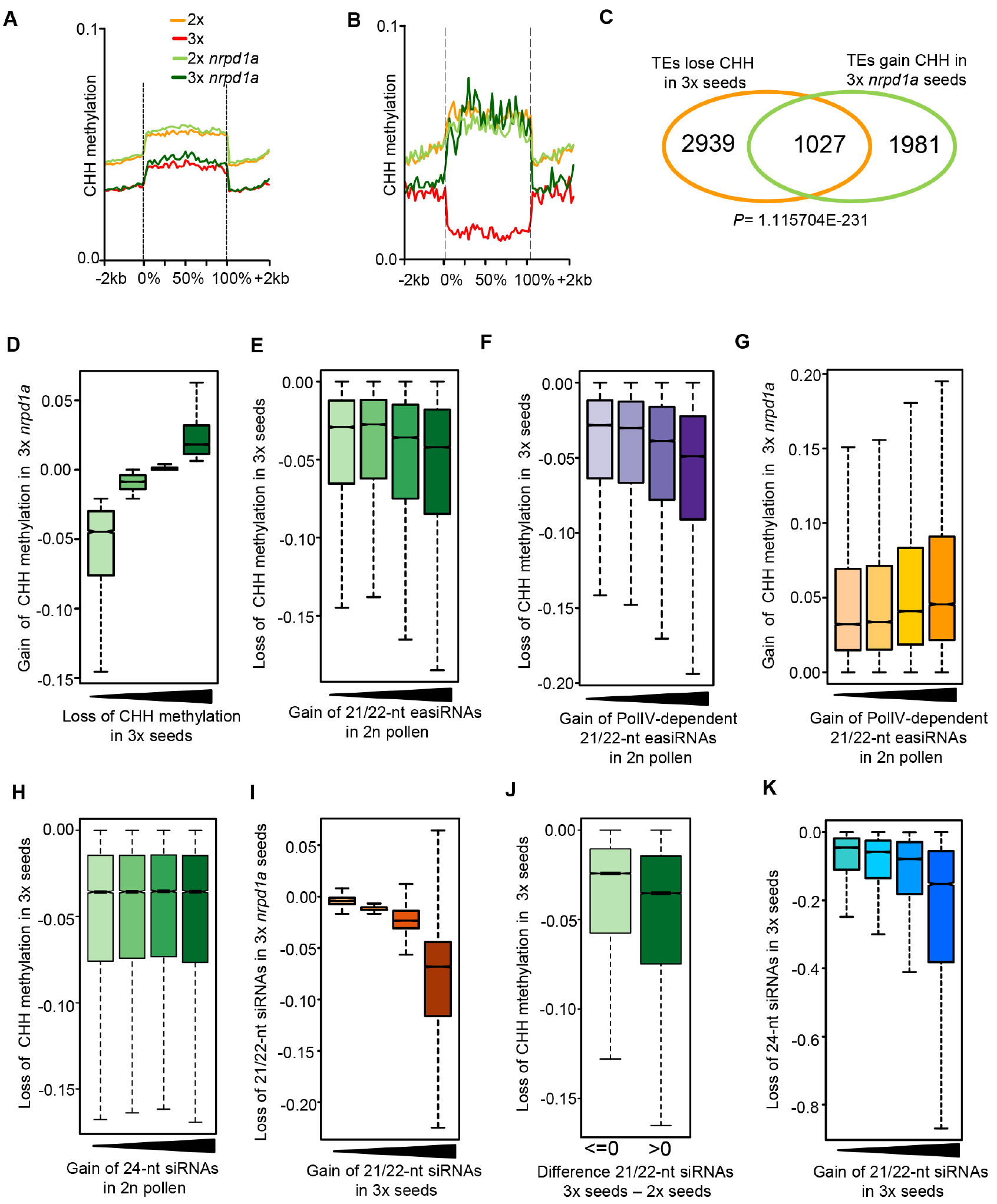
Triploid seeds have lower CHH methylation as a consequence of increased pollen-derived 21/22-nt easiRNAs. (**A**) CHH methylation metaplots over all TEs in the endosperm derived from 2×, 3×, 2× *nrpd1a*, and 3× *nrpd1a* seeds. (**B**) CHH methylation metaplots over TEs that lose CHH methylation in the endosperm of 3× seeds and restore CHH methylation in 3× *nrpd1a osd1* seeds. Color code as specified in A. (**C**) Venn-diagram showing overlap of TEs specified in **B**. Hypergeometric testing was used to test for significance of overlap. (**D-F**) Loss of CHH methylation in the endosperm of 3× seeds associates with increasing levels of CHH methylation in 3× *nrpd1a* seeds (**D**), increasing levels of 21/22-nt easiRNAs in 2n pollen (**E**), increasing levels of POL IV-dependent21/22-nt easiRNAs in 2n pollen (**F**). (**G**) Gain of CHH methylation in the endosperm of 3× *nrpd1a* seeds associates with increasing loss of 21/22-nt easiRNAs in 2n *nrpd1a* pollen. (**H**) Increasing levels of 24-nt easiRNAs in 2n pollen do not associate with CHH methylation differences in the endosperm of 3× seeds. (**I**) Increasing levels of 21/22-nt siRNAs in 3× seeds associate with increasing loss of 21/22-nt siRNAs in 3× *nrpd1a* seeds. (**J**) Increasing levels of 21/22-nt siRNAs in 3× seeds associate with loss of CHH methylation in 3× seeds. Plotted are differences of CHH methylation in the endosperm of 3× and 2× seeds at those 50 bps bins where differences in 21/22-nt siRNAs between 3× and 2× seeds are smaller or larger than zero. (**K**) Increasing levels of 21/22-nt siRNAs in 3× versus 2× seeds associate with increasing loss of 24-nt siRNAs in triploid seeds. (**D-I, K**) Plotted are CHH methylation or siRNA differences sorted by quantiles of 50-bp genome bins against differences of CHH methylation or siRNAs in the endosperm of indicated genotypes. Differences between first and last categories in **A-G** and **I-K** are significant (*P*<0.00001, Kolmogorov-Smirnov test).

We further challenged the hypothesis that 2n pollen contributes an increased dosage of 21/22-nt easiRNAs by testing whether increased levels of 21/22-nt siRNAs in 3× seeds correlate with loss of 21/22-nt siRNAs in 3× *nrpd1a* seeds (Fig. S7, Table S4). Loci with highest levels of 21/22-nt siRNAs in 3× seeds experienced the strongest loss of 21/22-nt siRNAs in 3× *nrpd1a* seeds (Fig. 3i), strongly supporting the idea that 2n pollen contributes an increased dosage of Pol IV-dependent easiRNAs that negatively interfere with CHH methylation in 3× seeds. Consistently, increased abundance of 21/22-nt siRNAs in 3× seeds correlated with increased loss of CHH methylation (Fig. 3j). Gain of 21/22-nt siRNAs in 3× seeds was associated with loss of 24-nt siRNAs (Fig. 3k), consistent with the loss of CHH methylation observed at loci targeted by 21/22-nt easiRNAs (Fig. 3j).

To address the question whether increased abundance of Pol IV-dependent easiRNAs in pollen associate with increased gene expression in the endosperm of 3× seeds, we generated transcriptome data of purified endosperm (Table S5). Increased abundance of Pol IV-dependent easiRNAs in pollen was associated with upregulated gene expression in the endosperm of 3× seeds (Fig. 4a), supporting the idea that increased abundance of easiRNAs negatively interferes with gene repression. Consistently, depletion of pollen-derived easiRNAs normalized gene expression in the endosperm of 3× seeds (Fig. 4b). Out of 57 PEGs with significantly increased expression in 3× seeds, 19 PEGs were at least 2-fold downregulated in 3× *nrpd1a* seeds (Fig. 4c^6,9^. Those 19 PEGs accumulated Pol IV-dependent easiRNAs at flanking regions (Fig. 4d), correlating with reduced CHH methylation in 3× seeds and partially restored CHH methylation in 3× *nrpd1a* seeds (Fig. 4e,f). Together weconclude that epigenetic changes at TEs are connected to changes of PEG expression.

**Fig. 4.**
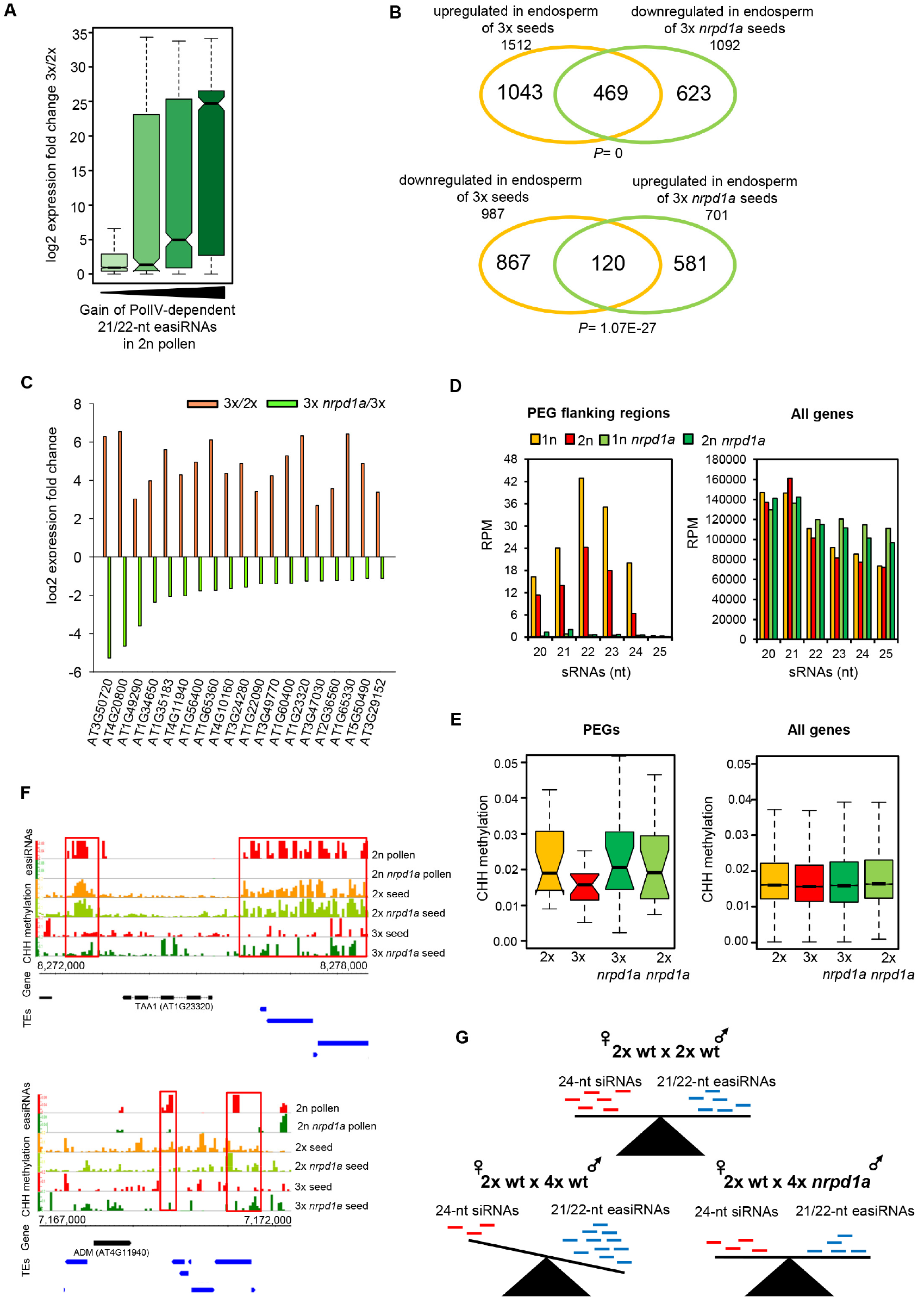
Pollen-derived 21/22-nt easiRNAs associate with gene expression changes in the endosperm of triploid seeds. (**A**) Increasing levels of 21/22-nt easiRNAs in 2n versus 2n *nrpd1a* pollen associate with upregulated gene expression in the endosperm of 3× seeds. Plotted are differences in 21/22-nt siRNAs in 2n and 2n *nrpd1a* pollen against log2-fold expression changes in the endosperm of 3× and 2× seeds. Differences between categories are significant (*P*<0.00001, Kolmogorov-Smirnov test). (**B**) Venn diagram showing overlap of deregulated genes in in the endosperm of 3× and 3× *nrpd1a* seeds compared to the corresponding 2× and 3× controls, respectively. Hypergeometric testing was used to test for significance of overlap. (**C**) Log2-fold change expression of PEGs in the endosperm of 3× and 3× *nrpd1a* compared to compared to the corresponding 2× and 3× controls, respectively. (**D**) sRNAs mapping to 2kb flanking regions of PEGs (specified in C) and flanking regions of all genes (left and right panels, respectively) in 1n and 2n wild-type and *nrpd1a* pollen. (**E**) Boxplots of CHH methylation in 2kb flanking regions of PEGs (specified in C) and all genes (left and right panels, respectively). (**F**) Representative PEGs accumulating 21/22-nt easiRNAs in pollen and experiencing changes in CHH methylation in the endosperm of 3× and 3× *nrpd1a* seeds. Red rectangles mark regions where easiRNA accumulation co-occurs with changes of CHH methylation. (**G**) Viable seed formation requires balanced populations of pollen-derived 21/22-nt easiRNAs and ovule-derived 24-nt siRNAs. Increased levels of 21/22-nt easiRNAs delivered from a higher ploidy pollen parent will reduce 24-nt siRNAs post-fertilization in the seed, resulting in unviable seed formation. *NRPD1a* deficient pollen restores viable seed formation by reducing pollen-derived easiRNAs.

Accordingly, we propose that Pol IV-dependent production of TE transcripts in gametic tissues is a genomic safeguard mechanism that sustains production of 21/22-nt easiRNAs of potentially harmful active TEs without Pol II transcription. Our data reveal a striking analogy of easiRNAs in establishing the triploid block with Piwi-interacting RNAs (piRNAs) in hybrid dysgenesis in flies ^23^. In both models, TE-derived sRNAs transmit epigenetic information transgenerationally, pointing to a conserved role of TE-derived sRNAs in assessing gamete compatibility. Pollen-delivered 21/22-nt easiRNAs could possibly target transcriptionally active TEs for degradation in the hypomethylated endosperm and thus prevent transposition ^24^. When different species hybridize, hybrid failure will result if pollen 21/22-nt easiRNAs do not recognize their maternal target TEs or, conversely, if paternal TEs are too sequence divergent to be recognized by maternal 24-nt siRNAs. Importantly, our data show that also inadequate dosage of parental siRNA populations can cause hybrid seed failure, revealing that the balanced dosage of maternal and paternal siRNA populations is essential for viable seed formation (Fig. 4e). One possible scenario is that increased dosage of 21/22-nt easiRNAs causes these siRNAs to target scaffolding transcripts produced by either Pol II or Pol V and thus negatively interfere with DNA methylation establishment, as previously proposed ^12^.

The triploid block has been a mystery to geneticists and breeders, formalized as the “endosperm balance number’ hypothesis more than 35 years ago ^1^. Our discovery that paternal easiRNAs form the genetic basis of dosage sensitivity and establish the triploid block provides the means for targeted strategies bypassing this hybridization barrier.

## Methods

### Plant material and growth conditions

The mutants used in this study were described previously as *ago1-36* (Salk_087076; ^25^, *ago2-1* (Salk_003380; ^26^, *ago6-2* (Salk_031553; ^27^, *rdr2-2* (Salk_059661; ^20^, *rdr6-15* (Sail_617H07; ^28^, *nrpd1a-3* (Salk_128428; ^29^, *nrpd1b-12* (Salk_033852; ^30^, *dcl2-1* (Salk_064627; ^31^, *dcl3-1* (Salk_005512; ^31^, *dcl4-2* ^31^. The *osd1-1* mutant ^5^ was kindly provided by Raphael Mercier. Being originally identified in the Nossen background, the mutant was introgressed into Col by repeated backcrosses over five generations. Tetraploid *nrpd1a-3* plants were generated using colchicine treatment as previously described ^22^. Plants were grown in a growth cabinet under long day photoperiods (16 hr light and 8 hr dark) at 22°C. After 10 days, seedlings were transferred to soil and plants were grown in a growth chamber at 60% humidity and daily cycles of 16 hr light at 22°C and 8 hr darkness at 18°C. For all crosses, designated female plants were emasculated and the pistils were hand-pollinated 2 days after emasculation.

### Germination analysis

Seeds were surface sterilized in a container using chlorine gas (10 ml hydrochloric acid plus 50 ml sodium hypochlorite) and incubated for up to 3 hr. To determine germination frequency, seeds were plated on 1/2 MS media containing 1% sucrose, stratified at 4°C for 2 days in the dark and grown in a growth cabinet under long day photoperiods (16 hr light and 8 hr dark) at 22°C for 10 days.

### Microscopy

Seeds were fixed and embedded with Technovit 7100 (Heraeus, Germany) as described ^32^. Five-micrometer sections were prepared with an HM 355 S microtome (Microm, Germany) using glass knives. Sections were stained for 1 min with 0.1% toluidine blue and washed three times with destilled water. Microscopy was performed using a DMI 4000B microscope with DIC optics (Leica, Germany). Images were captured using a DFC360 FX camera (Leica) and processed using Photoshop CS5 (Adobe, San Jose, California).

### Pollen normalization and Northern blotting

10 ul of total pollen extracts of different genotypes was counted under a microscope. Based on the determined pollen density different volumes of the extracts were used for subsequent downstream RNA extraction. Northern blotting was done as previously described ^10^.

### RNA sequencing

For RNA sequencing, endosperm from seeds derived from crosses L*er* × Col-0, L*er* × *osd1*, L*er* × *osd1 nrpd1a* and L*er* × *nrpd1a* were dissected in biological duplicates following previously described procedures ^22^. RNA was extracted following a modified protocol for the RNAqueous kit (Ambion, Life Technologies). RNA was purified by Qiagen RNeasy Plant Mini Kit (Qiagen, Hilden, Germany) after residual DNA was removed by adding 2 uL DNaseI (Thermo-Scientific, Waltham, USA). Libraries were prepared using the TruSeq RNA Library Prep Kit v2 (Illumina, San Diego, USA) and sequenced at the SciLife Laboratory (Uppsala, Sweden) on an Illumina HiSeq2000 in 50-bp single-end mode.

### Small RNA sequencing

Pure mature pollen samples from inflorescences of approximately 500 plants were collected as previously described ^33^ with minor modifications. Flowers were harvested in a beaker, covered with 9% sucrose solution and shaken vigorously for 5 min to release pollen grains into the solution. The subsequent centrifugation- and filtering-steps were carried out in 9% sucrose solution. The obtained pollen pellet was frozen in liquid nitrogen and stored at −70 °C. Total RNA was extracted using Trizol reagent (Ambion/ Life Technologies, USA) and glass beads (1.25-1.55 mm; Carl Roth) and 10 μg of total RNA were run on a 14% TBE UREA Polyacrylamide gel for size selection. Gel slices containing RNA in the range of 17- to 25-nt were purified following the Illumina TrueSeq small RNA protocol for gel extraction. Libraries were constructed using the Illumina TruSeq Small RNA library preparation kit (RS-200-0012) according to the manufacturer’s instructions.

To generate sRNA libraries from seeds, we crossed male sterile *pistillata* maternal plants (in Ler background ^34^) with pollen from Col-0, 4× Col-0, *nrpd1a*, and 4× *nrpd1a*. For each genotype 1000 seeds at 6-7 DAP were collected in duplicates in RNA later (Sigma-Aldrich) and homogenized (Silamat S5) using glass beads. Total RNAs of seeds were extracted using mirVana miRNA isolation kit (Ambion/Life Technologies) and sRNAs were isolated by FDF-PAGE ^35^. SRNA libraries were generated using the NEBNext^®^ Multiplex Small RNA Library Prep Set for Illumina. Libraries were sequenced at the SciLife Laboratory (Uppsala, Sweden) on an Illumina HiSeq2000 in 50-bp single-end fashion.

### Bisulfite sequencing

To generate bisulfite libraries we dissected endosperm from seeds derived from crosses L*er* × Col-0, L*er* × *osd1*, L*er* × *osd1 nrpd1a* and L*er* × *nrpd1a* in biological duplicates following previously described procedures ^22^. DNA purification and library preparation were done as described in ^36^. Quality of isolated endosperm was calculated based on Col and Ler SNPs (Table S2).

### mRNA sequencing data analysis

For each replicate, 50 bp long reads were mapped to the Arabidopsis (TAIR10) genome, masked for rRNA genes, using TopHat v2.1.0 (Trapnell et al, 2009) (parameters adjusted as -g 1 -a 10 -i 40 -I 5000 -F 0 -r 130). Gene and TE expression was normalized to reads per kilobase per million mapped reads (RPKM) using GFOLD ^37^. Expression level for each condition was calculating using the mean of the expression values in both replicates. Differentially regulated genes and transposable elements across the two replicates were detected using the rank product method as implemented in the Bioconductor RankProd Package ^38^. The test was run with 100 permutations and gene selection was corrected for multiple comparison errors using a pfp (percentage of false prediction) < 0.05.

### Small RNA sequencing data analysis

Adapters were removed from the 50 bp long single-end sRNA reads in each library. The resulting18-30 bp long reads were mapped to a Col genome (TAIR10) masked for Ler SNPs using bowtie (-v 2 −best). All reads mapping to chloroplast and mitochondria and to structural noncoding RNAs (tRNAs, snRNAs, rRNAs, or snoRNAs) were removed. Mapped reads from both replicates were pooled together, sorted in two categories (21-22-nt and 24-nt long) and remapped to the same reference masked genome mentioned above using ShorStack (--mismatches 2 --mmap f) ^39^ in order to improve the localization of sRNAs mapping to multiple genomic locations. We normalized the alignments by converting coverage values to reads per million mapped reads (RPM) values. The sRNA mapping profiles were visualized with bedGraph files based in 50 bp bins. Metagene plots over TEs were constructed between −2 kb and + 2 kb by calculating mean levels of sRNA (RPM) in 100 bp bins in the flanks of the TEs and in 40 equally long bins between the transcriptional start and stop.

### DNA methylation analysis

For each mutant, the 125 bp reads from the Illumina BS sequence libraries from the two biological replicates were merged and trimmed to 100 bp by cutting 5 bp from the start and 20 bp from the end of each read. After this first trimming, each read was split in two 50 bp long reads in order to improve mapping efficiency.

Reads were mapped to the TAIR10 Arabidopsis genome using the Bismark read mapper ^40^ allowing up to one mismatch per read. Duplicated reads (aligning to the same genomic position) were eliminated before calculating methylation levels. Methylation levels for each condition were calculated as the mean of the two replicates. Cytosine conversion efficiency was estimated as the percentage of CHH methylation in the chloroplast. Cytosine methylation was visualized separately for CG, CHG and CHH cytosine contexts with bedGraph files representing average methylation values in 50 bp bins across the genome. Differential analysis on the levels of CHH methylation in TEs between conditions considered the cytosine methylation reports from both replicates and was performed using linear modelling (p-value < 0.05) as implemented in the R package Limma ^41^.

## Acknowledgements

We thank Rob Martienssen and Filipe Borges for critical comments on the manuscript. This research was supported by a European Research Council Starting Independent Researcher grant (to C.K.), a grant from the Swedish Science Foundation (to C.K.) and a grant from the Knut and Alice Wallenberg Foundation (to C.K.). G.M. was supported by a Marie Curie IOF Postdoctoral Fellowship (PIOF-GA-2012-330069). Sequencing was performed by the SNP&SEQ Technology Platform, Science for Life Laboratory at Uppsala University, a national infrastructure supported by the Swedish Research Council (VRRFI) and the Knut and Alice Wallenberg Foundation.

## Author Contributions

GM, PW, RKS and CK designed the experiments. GM, PW, ZW, J M-R, CDeF performed the experiments and generated the data. GM, JS-G and LLC carried out the bioinformatic analysis. GM, PW, JS-G, LLC, RKS and CK analyzed the data. GM, PW and CK wrote the manuscript.

## References

1. Johnston, S., Nijs, T., Peloquin, S. & Hanneman, R. The significance of genic balance to endosperm development in interspecific crosses. Theor. Appl. Genetics 57, 5–9 (1980).

2. Schatlowski, N. & Köhler, C. Tearing down barriers: understanding the molecular mechanisms of interploidy hybridizations. J Exp Bot 63, 6059–67 (2012).

3. Scott, R.J., Spielman, M., Bailey, J. & Dickinson, H.G. Parent-of-origin effects on seed development in Arabidopsis thaliana. Development 125, 3329–3341 (1998).

4. Lu, J., Zhang, C., Baulcombe, D.C. & Chen, Z.J. Maternal siRNAs as regulators of parental genome imbalance and gene expression in endosperm of Arabidopsis seeds. Proc Natl Acad Sci U S A 109, 5529–5534 (2012).

5. d'Erfurth, I. et al. Turning meiosis into mitosis. PLoS Biol 7, e1000124 (2009).

6. Kradolfer, D., Wolff, P., Jiang, H., Siretskiy, A. & Köhler, C. An imprinted gene underlies postzygotic reproductive isolation in Arabidopsis thaliana. Dev Cell 26, 525–535 (2013).

7. Law, J.A. & Jacobsen, S.E. Establishing, maintaining and modifying DNA methylation patterns in plants and animals. Nat Rev Genet 11, 204–220 (2010).

8. Havecker, E.R. et al. The Arabidopsis RNA-directed DNA methylation argonautes functionally diverge based on their expression and interaction with target loci. Plant Cell 22, 321–334 (2010).

9. Wolff, P., Jiang, H., Wang, G., Santos-Gonzalez, J. & Kohler, C. Paternally expressed imprinted genes establish postzygotic hybridization barriers in Arabidopsis thaliana. Elife 4, doi:10.7554/eLife.1 (2015).

10. McCue, A.D., Nuthikattu, S., Reeder, S.H. & Slotkin, R.K. Gene expression and stress response mediated by the epigenetic regulation of a transposable element small RNA. PLoS Genet 8, e1002474 (2012).

11. Nuthikattu, S. et al. The initiation of epigenetic silencing of active transposable elements is triggered by RDR6 and 21-22 nucleotide small interfering RNAs. Plant Physiol 162, 116–131 (2013).

12. Creasey, K.M. et al. miRNAs trigger widespread epigenetically activated siRNAs from transposons in Arabidopsis. Nature 508, 411–415 (2014).

13. McCue, A.D. et al. ARGONAUTE 6 bridges transposable element mRNA-derived siRNAs to the establishment of DNA methylation. EMBO J 34, 20–35 (2015).

14. Martinez, G., Panda, K., Köhler, C. & Slotkin, R.K. Silencing in sperm cells is directed by RNA movement from the surrounding nurse cell. Nature Plants 2, 16030 (2016).

15. Slotkin, R.K. et al. Epigenetic reprogramming and small RNA silencing of transposable elements in pollen. Cell 136, 461–472 (2009).

16. Li, S. et al. Detection of Pol IV/RDR2-dependent transcripts at the genomic scale in Arabidopsis reveals features and regulation of siRNA biogenesis. Genome Res 25, 235–245 (2015).

17. Zhai, J. et al. A One Precursor One siRNA Model for Pol IV-Dependent siRNA Biogenesis. Cell 163, 445–455 (2015).

18. Wei, W. et al. A role for small RNAs in DNA double-strand break repair. Cell 149, 101–112 (2012).

19. Gasciolli, V., Mallory, A.C., Bartel, D.P. & Vaucheret, H. Partially redundant functions of Arabidopsis DICER-like enzymes and a role for DCL4 in producing trans-acting siRNAs. Curr Biol 15, 1494–1500 (2005).

20. Vazquez, F. et al. Endogenous trans-acting siRNAs regulate the accumulation of Arabidopsis mRNAs. Mol Cell 16, 69–79 (2004).

21. Peragine, A., Yoshikawa, M., Wu, G., Albrecht, H.L. & Poethig, R.S. SGS3 and SGS2/SDE1/RDR6 are required for juvenile development and the production of trans-acting siRNAs in Arabidopsis. Genes Dev 18, 2368–2379 (2004).

22. Schatlowski, N. et al. Hypomethylated pollen bypasses the interploidy hybridization barrier in Arabidopsis. Plant Cell 26, 3556–3568 (2014).

23. Brennecke, J. et al. An epigenetic role for maternally inherited piRNAs in transposon silencing. Science 322, 1387–1392 (2008).

24. Martienssen, R.A. Heterochromatin, small RNA and post-fertilization dysgenesis in allopolyploid and interploid hybrids of Arabidopsis. New Phytol 186, 46–53 (2010).

25. Baumberger, N. & Baulcombe, D.C. Arabidopsis ARGONAUTE1 is an RNA Slicer that selectively recruits microRNAs and short interfering RNAs. Proc Natl Acad Sci U S A 102, 11928–11933 (2005).

26. Lobbes, D., Rallapalli, G., Schmidt, D.D., Martin, C. & Clarke, J. SERRATE: a new player on the plant microRNA scene. EMBO Rep 7, 1052–1058 (2006).

27. Zheng, X., Zhu, J., Kapoor, A. & Zhu, J.K. Role of Arabidopsis AGO6 in siRNA accumulation, DNA methylation and transcriptional gene silencing. EMBO J 26, 1691–1701 (2007).

28. Allen, E. et al. Evolution of microRNA genes by inverted duplication of target gene sequences in Arabidopsis thaliana. Nat Genet 36, 1282–1290 (2004).

29. Herr, A.J., Jensen, M.B., Dalmay, T. & Baulcombe, D.C. RNA polymerase IV directs silencing of endogenous DNA. Science 308, 118–120 (2005).

30. Pontier, D. et al. Reinforcement of silencing at transposons and highly repeated sequences requires the concerted action of two distinct RNA polymerases IV in Arabidopsis. Genes Dev 19, 2030–2040 (2005).

31. Xie, Z. et al. Genetic and functional diversification of small RNA pathways in plants. PLoS Biol 2 (2004).

32. Hehenberger, E., Kradolfer, D. & Köhler, C. Endosperm cellularization defines an important developmental transition for embryo development. Development 139, 2031–2039 (2012).

33. Schoft, V.K. et al. SYBR Green-activated sorting of Arabidopsis pollen nuclei based on different DNA/RNA content. Plant Reprod 28, 61–72 (2015).

34. Goto, K. & Meyerowitz, E.M. Function and regulation of the Arabidopsis floral homeotic gene pistillata. Genes Dev 8, 1548–1560 (1994).

35. Harris, C.J., Molnar, A., Muller, S.Y. & Baulcombe, D.C. FDF-PAGE: a powerful technique revealing previously undetected small RNAs sequestered by complementary transcripts. Nucleic Acids Res 43, 7590–7599 (2015).

36. Moreno-Romero, J., Jiang, H., Santos-Gonzalez, J. & Kohler, C. Parental epigenetic asymmetry of PRC2-mediated histone modifications in the Arabidopsis endosperm. EMBO J 35, 1298–1311 (2016).

37. Feng, J. et al.. GFOLD: a generalized fold change for ranking differentially expressed genes from RNA-seq data. Bioinformatics 28, 2782–2788 (2012).

38. Hong, F. et al. RankProd: a bioconductor package for detecting differentially expressed genes in meta-analysis. Bioinformatics 22, 2825–2827 (2006).

39. Johnson, N.R., Yeoh, J.M., Coruh, C. & Axtell, M.J. Improved Placement of Multi-mapping Small RNAs. G3 (Bethesda) 6, 2103–2111 (2016).

40. Krueger, F. & Andrews, S.R. Bismark: a flexible aligner and methylation caller for Bisulfite-Seq applications. Bioinformatics 27, 1571–1572 (2011).

41. Ritchie, M.E. et al. Limma powers differential expression analyses for RNA-sequencing and microarray studies. Nucleic Acids Res 43, e47 (2015).

